# Discrimination of Normal from Slow-Aging Mice by Plasma Metabolomic and Proteomic Features

**DOI:** 10.1101/2025.05.11.651908

**Authors:** Bretton Badenoch, Oliver Fiehn, Noa Rappaport, Pranjal Srivastava, Kengo Watanabe, Sriram Chandrasekaran, Richard A. Miller

## Abstract

Tests that can predict whether a drug is likely to extend mouse lifespan could speed up the search for anti-aging drugs. We have applied a machine learning algorithm, XGBoost regression, to seek sets of plasma metabolites that can discriminate control mice from mice treated with an anti-aging diet (caloric restriction) or any of four anti-aging drugs. When the model is trained on any four of these five interventions, it predicts significantly higher lifespan extension in mice exposed to the intervention which was not included in the training set. Plasma peptide data sets also succeed at this task. Models trained on drug-treated normal mice also discriminate long-lived mutant mice from their respective controls, and models trained on males can discriminate drug-treated from control females. Triglycerides are over-represented among the most influential features in the regression models. Triglycerides with longer fatty acid chains tend to be higher in the slow-aging mice, while triglycerides with shorter fatty acid chains tend to decrease. Plasma metabolite patterns may help to select the most promising anti-aging drugs in mice or in humans, and may give new leads into physiological and enzymatic targets relevant to discovery of new anti-aging drugs.

## Introduction

Aging is associated with higher risks of most major contributors to mortality in developed countries, such as cancer, cardiovascular disease and neurodegeneration (1). With the aim of postponing aging and age-associated diseases, the National Institute on Aging has for 20 years supported a systematic search for anti-aging drugs (2). Eleven distinct agents have produced, so far, significant increases in lifespan in one or both sexes, including acarbose (Aca), rapamycin (Rapa), 17-a-estradiol (17aE2), and canagliflozin (Cana) (3–6). Direct tests of proposed new anti-aging drugs are slow and expensive, however, because lifespan studies in mice require a minimum of 36 months to complete.

Multi-omics analyses, particularly metabolomics and proteomics, have yielded valuable insights into how various anti-aging interventions alter molecular pathways in mice. For instance, partial chemical reprogramming of murine fibroblasts using a small-molecule cocktail significantly enhanced oxidative phosphorylation and reduced aging-related metabolites (7). In another study, treatment with 17aE2 was found to ameliorate sarcopenia in male mice, evidenced by changes in muscle metabolite levels, chiefly amino acids (8). Similarly, Aca and Rapa modulated the cardiac and plasma lipidomes of mice (9).

Moreover, proteomic and metabolomic profiling of long-lived growth hormone-releasing hormone knock-out (GHRH-KO) mice emphasized the importance of enhanced mitochondrial function and altered amino acid metabolism in lifespan extension (10). Collectively, these studies suggest that there are detectable patterns of metabolic change in mice treated with anti-aging compounds, but it is unknown to what degree these interventions overlap.

Our current study makes use of metabolomic and proteomic (peptide-level) data obtained from plasma of young adult (i.e. 12 month old), genetically heterogeneous (2, 11) mice that had been exposed to an anti-aging drug or to a calorie-restricted (CR) diet from age 4 months. The data set included information on over 12,000 different metabolites and over 17,000 identified peptides. We used a machine learning approach (XGBoost regression) (12) For each mouse, the percentage lifespan increase was taken to be the median lifespan increase in published data for the intervention in question, and as zero percent for untreated control mice. Feature abundance was regressed on this estimated lifespan increase. Estimated lifespan increase for each individual mouse was then calculated based on the abundance of features using the trained regression model. This approach allowed us to elucidate whether metabolomics or proteomics data, or the combination, could serve to make reliable predictions. We also used a “novel intervention test” method, in which the regression model was trained using only four of the five available groups of treated mice, and then used to calculate the estimated lifespan increase for each mouse in the group that was not included in the training procedure, to simulate a situation in which plasma samples were available for mice treated with candidate anti-aging drugs whose effect on lifespan was unknown. We also noted the specific features that contribute most strongly to be performance of each model, with particular attention to changes in the fatty acid constituents of specific triglycerides.

## Methods

### Mouse Samples

UM-HET3 mice were weaned at 19 – 21 days of age and housed at 3 males/cage or 4 females/cage, using the same conditions developed for the Interventions Testing Program (2). At four months of age, mice in the CR group were given 80% of the amount of chow consumed by age- and sex-match control mice, reduced to 60% at five months of age. CR mice were fed once/day, between 8 – 9 am. All other mice had ad libitum access to food. UM-HET3 mice in the other intervention groups began drug treatment at 4 months of age. Mice were euthanized at 12 months of age, between 8 am and 11 am, using CO2 asphyxiation. Mice lost consciousness within 10 seconds and were removed for closed-chest cardiac puncture as soon as breathing stopped, typically within 30 seconds. Blood samples were collected using heparin-coated syringes, and plasma removed by centrifugation was then aliquoted and stored at −80°C until evaluation.

Production of GHRKO and Snell dwarf mice used the breeding schemes described previously (13, 14). GHRKO mice and their non-mutant littermates were on the C57BL/6 background, and Snell dwarf and their non-mutant littermates were on a segregating stock including 25% C3H/HeJ and 75% DW/J background genes.

### Analysis of Metabolites in Plasma

Plasma metabolites were acquired by five different mass spectrometry techniques: four liquid chromatography-accurate mass spectrometry methods (LC-MS) and one gas chromatography-mass spectrometry assay (GC-MS). Details are given in (15). Briefly, 20 ul plasma samples were extracted by 1 ml methanol/water/MTBE (16), yielding a lipophilic phase and a hydrophilic phase. Extraction phases were separated, dried, and prepared for LC-MS and GC-MS assays. Data was acquired by hydrophilic interaction chromatography (HILIC)-Orbitrap QE HF mass spectrometry under positive and negative mode electrospray ionization for polar metabolites (17), plus positive and negative mode electrospray ionization for lipids using C18 reversed phase chromatography (RPLC)-Orbitrap QE HF mass spectrometry (18), plus positive mode electron ionization GC-MS using a nominal mass time of flight mass spectrometer (Leco Pegasus IV GC-TOF MS). Data from LC-MS assays was processed in MS-DIAL 4.90 with compound annotations using NIST20 and MassBank.us mass spectral libraries. GC-TOF MS data was processed in ChromaTOF 4.0 software with compound annotations in the BinBase database (19). Data was normalized using random forest machine learning SERRF software by matching to quality control pool samples (20). Annotated metabolites (at Metabolomics Society Initiative levels 1-3, MSI (21)) were filtered to have signal/noise ratios s/n>3, and unknown metabolites were filtered at s/n>10 in comparison to method blank negative controls for local noise estimations.

### Analysis of Peptides in Plasma

Plasma samples from 284 mice were processed for data-independent acquisition-parallel accumulation and serial fragmentation (DIA-PASEF) analysis as follows. Frozen plasma aliquots were thawed at 4 °C for 3 h followed by a hard spin to pellet insoluble particles (5 min, 10000g, 4 °C). Protein concentration was determined using the BCA assay (Pierce, Cat# 23227). Samples were normalized by aliquoting 400 μg in phosphate-buffered saline (PBS) into a final volume of 37.5 μL. Denaturation was performed by adding a 12.5 μL lysis buffer (200 mM triethylammonium bicarbonate (TEAB), 20% SDS) to a final concentration of 5% SDS and heating for 5 min at 90 °C with 800 rpm shaking. After denaturation, samples were reduced with 5 mM TCEP (Sigma, USA Cat# 4706) for 15 min at 55 °C and then alkylated with 10 mM iodoacetamide (Millipore-Sigma, USA, Cat# 407710) for 10 min in the dark at room temperature, both with 800 rpm shaking. Samples were acidified with neat orthophosphoric acid to a final concentration of 2.7% (v/v) before applying the sample to S-Traps (ProtiFi, USA). The S-trap buffer (100 mM TEAB pH 8.5/90% methanol) was added at a 1:7 ratio prior to loading on the S-trap 96 well-plate format by a 1 min spin at 4000g. Once loaded, samples were washed six times with 400 μL of S-trap buffer and spun at 4000g for 5 min to ensure complete dryness.

Tryptic digestion was performed at 37 °C with 125 μL of trypsin (Promega, USA Cat# V511X) in a digestion buffer (100 mM TEAB pH 8.5) at a 1:25 ratio. After the first hour of incubation at 37 °C, an additional 75 μL of digestion buffer was added to prevent the drying out of the samples. After overnight incubation at 37 °C, the digested peptides were eluted with 80 μL of digestion buffer and centrifugation at 4000g for 1 min, and then with 80 μL of 50% ACN in a digestion buffer and centrifugation at 4000g for 1 min. The final eluate was ∼250 μL. Quantification of the digested peptides was performed by the fluorescamine fluorescent peptide assay (22) (Pierce, USA, Cat# 23290) before drying down to completion.

All peptide samples were spiked in with iRT standard peptides (Biognosys AG, Schlieren, Switzerland) and applied to MS analysis using a Vanquish Neo HPLC system (Thermo-Fisher Scientific, USA), configured in microflow mode and coupled to a timsTOF PRO mass spectrometer (Bruker, USA). The Vanquish Neo HPLC system was operated with 99.9% water, 0.1% formic acid/Milli-Q water (v/v, Buffer A), and 99.9% ACN, 0.1% formic acid (v/v, Buffer B).

For VIP-HESI measurements, the microflow workflow was used with Vanquish Neo. Peptides were trapped on a 50 × 1 mm ID trap cartridge Chrom XP C18, 3 μm (Thermo-Fisher Scientific, USA) at 50 μL/min and separated on a C18, 15 cm × 1 mm × 1.7 μm Kinetix column (Phenomenex, USA) at 40 μL/min using a 45 min linear gradient. The VIP-HESI source was equipped with a 50 μm electrode probe (Bruker, USA), and the parameters were as follows: 4000 V capillary voltage, 3.0 L/min dry gas, and temperature 200 °C, probe gas flow 3.0 L/min and temperature 100 °C. The VIP-HESI source parameter settings were optimized for spray stability over extended periods of time using the background signal. Neat mouse plasma sample injections were acquired using Bruker timsTOF preformed DIA-PASEF mode schema covering the m/z range of 400–1200 and 1/K0 range 0.6 to 1.42 in 32 × 25 Da windows with a mass overlap of 1 Da, resulting in a total cycle time of 1.8 s. 40 μg of mouse plasma sample were measured and the high sensitivity mode was enabled in the tims control acquisition software of the mass spectrometer (23).

### Proteomics Data Analysis

Spectral Assay Library Quality Assessment Using DIALib-QC and data processing using Spectronaut. Comprehensive mouse plasma spectral assay libraries were was assessed for their quality using DIALib-QC (v1.2) (24). DIALib-QC evaluates 57 parameters of compliance and provides a detailed report of the library’s complexity, characteristics, modifications, completeness, and correctness. In the DIALib-QC assessment report, there were no problem assays found for both libraries, which were used as it is for DIA-MS analysis.

Spectronaut (Biognosys, Switzerland) DIA software tools were used in this study to process data from the mouse plasma samples. Quantification and DIA processing of PepCalMix and control UM-HET mouse plasma samples were performed using Spectronaut DIA software (version 16.0.220606.53000 (Biognosys, Switzerland). For the nonlinear iRT calibration strategy, a dynamic window was used for both mass tolerance (MS1 and MS2), and to set up the extracted ion chromatogram (XIC) retention time (RT) window. Preprocessing of MS1 and MS2 calibration strategies was enabled. Decoy assays were dynamically generated using the scrambled decoy method with a set size of 0.1 as a fraction of the library size input. The identification was performed using the kernel density estimator with precursor and protein identification results filtered with a q-value of <0.01. For quantification, MS2 ion peak areas of quantified peptides were averaged to estimate the protein peak areas. Additional parameter settings were used as the default.

For PepCalMix sample data processing, the PepCalMix library was directly used, and the identification was performed using a normal distribution density estimator with precursor and protein identification results filtered with a q-value of <0.01. For quantification, the Top N ranking order setting was disabled to include all 20 peptides to estimate PepCalMix protein quantity. In the linearity experiment, the analysis for all seven different sample amounts was conducted together for each source. Similarly, for the impact of flow rates analysis, all sample amounts with replicates were analyzed together. For the direct comparison of PepCalMix peptide abundances between different ion sources, joint processing of the respective measurements was performed.

Data analysis of 284 plasma samples was performed using Spectronaut DIA software (version 17.0.221202.55965 (Biognosys, Switzerland) using both a mouse library (as described above) and directDIA (library-free mode) to increase the proteome coverage and reduce the sparsity in the combined data matrix. For directDIA workflow, the database and parameter settings were kept the same as described above. Default settings were used without global normalization enabled. Trypsin specificity was set to two missed cleavages and a false discovery rate of 1% on both peptide and protein were used. Data filtering was set to q-value.

### Datasets

The datasets used in this paper include plasma metabolomics and plasma peptides. Two versions of the metabolomic dataset were used. The AM data (“all metabolites”) included all the metabolites detected, while the FM (“filtered metabolite”) dataset included only structurally annotated metabolites with MSI = 1, 2, or 3. Similarly, we used two peptide datasets. The AP (“all peptide”) group contained data on each peptide detected, while the FP (“filtered peptide) dataset was created by retaining only peptides that had no missing data among the set of control mice. All four datasets were Log2 transformed. These datasets were also combined to create the AMP dataset (AM + AP) and FMP dataset (FM + FP). **Table 1** shows the number of features within each of these six datasets.

**Table 1:** Six data sets used.

| **Dataset** | **Number Of Features** | **Number of Male Mice** |
| --- | --- | --- |
| AM | 12,320 | 148 |
| FM | 1,051 | 148 |
| AP | 17,628 | 151 |
| FP | 6,651 | 151 |
| AMP | 29,949 | 127 |
| FMP | 7,701 | 127 |

The number of male mice (including control and treated groups) that contributed data to these six sets is shown in the table. All of the data for training were derived from UM-HET3 mice at 12 months of age. The rationale for focusing on models trained on data from males is presented in the text.

### Lifespan Estimation for Groups of Treated Mice

We utilized a machine learning approach, XGBoost Regression (12), to estimate the median percentage increase in lifespan associated with various treatments in mice, independently for each of the six available datasets shown in Table 1. XGBoost was selected due to its ability to handle large, complex datasets with high dimensionality and its effectiveness in identifying non-linear relationships between features and outcomes. For each feature (peptide or metabolite abundance), the dependent variable, median percentage increase in lifespan, was provided based on values in the published literature. **Table 2** shows, for each intervention, the published percent lifespan increase and the reference from which the increase was taken. The data are all based on studies of male UM-HET3 mice. Estimates for CR are from the Jackson Laboratory colony and estimates for the other four treatments are derived from the three ITP laboratories, i.e., pooled from Jackson Laboratory, University of Michigan, and University of Texas Health Science Center at San Antonio.

**Table 2:**
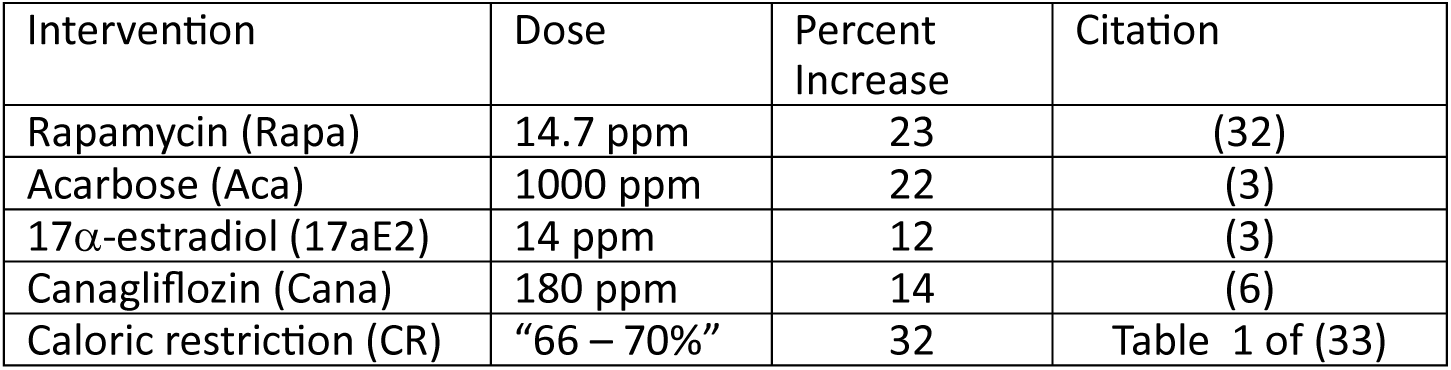
Published percent lifespan increase for male mice for interventions used in this study.

### XGBoost Regression Models

Data preprocessing involved excluding non-informative and identifier columns, retaining only numerical features relevant for modeling. We employed the Extreme Gradient Boosting (XGBoost) algorithm to model the relationship between the biological features and the median lifespan increase (12). The version information for all packages can be found for each module in the git repository (https://github.com/BrettonB/AgingDrugOmics/tree/main/Package_Info). To assess the model’s predictive performance, we initially implemented a 10-fold cross-validation strategy. Each dataset was randomly partitioned into ten equal subsets. In each iteration, one subset was used as the test set while the remaining nine subsets constituted the training set. This process was repeated across multiple iterations with different random seeds to ensure the stability of the results. Predictions from all folds and iterations were aggregated to compute median values for each treatment group within each dataset. Statistical significance between treatment groups and controls was assessed using independent Student’s t-tests.

### XGBoost Novel Intervention Test (NIT)

For this set of tests, XGBoost models were trained on the contrast between untreated controls and a “treated” group consisting of four of the five interventions (CR diet and four drugs, as in Table 2). The model was then used to estimate the predicted percent increase in median lifespan for each mouse in the treatment group that had been omitted from the training set. This procedure was then reiterated for each of the five varieties of intervention.

### SHAP Plots

SHAP values (SHapley Additive exPlanations) were calculated using the Shap package in Python for a model trained with all interventions. Features were then rank-ordered by SHAP statistic (25) to develop a list of the features with greatest influence on the estimated percent lifespan increase. The SHAP scores for each of the high-ranked features were then displayed as a log2-fold change graphic. The fold change was calculated by averaging the values of the mice within one intervention group, and then dividing by the age-matched control. Then the log2 of this value was displayed to visualize more easily increases and decreases.

### Statistics For Contrasts of Treated Group to Untreated Control Mice

The Student’s 2-tailed t-test was performed comparing the control group measurements to the novel intervention measurements.

### Code Availability

All custom scripts and workflows used in this study are publicly available at the following GitHub repository: https://github.com/BrettonB/AgingDrugOmics. Any additional code or analysis tools will be archived and assigned a DOI via Zenodo, and these details will be updated upon final acceptance of the manuscript. The plasma metabolite visualization browser tool is available at https://bretton-badenoch.shinyapps.io/Plasma-Anti-Aging-Mouse-Metabolites/.

### Data Availability

The metabolomic and peptide plasma LC–MS datasets analyzed during this study will be made publicly accessible via Zenodo (https://zenodo.org/). A permanent DOI and direct link to the datasets will be provided once the manuscript is accepted for publication. In the meantime, researchers seeking early access to the data may contact the corresponding author.

## Results

### XGBoost regression models trained on published percent lifespan increase for each treatment group

In initial studies, we compared the ability of several varieties of machine learning (ML) algorithms to distinguish between control mice and mice that had been exposed to anti-aging interventions (“treated” mice). The treated mouse group included mice exposed to either rapamycin (Rapa, 14.7 ppm), acarbose (Aca, 1000 ppm), 17α-estradiol (17aE2, 14 ppm), or canagliflozin (Cana, 180 ppm), or which had been exposed to a calorie restricted (CR) diet, adjusted to 60% of food consumed by age-matched mice. All interventions were initiated at 4 months of age, and plasma samples were taken from 12-month-old mice. **Table 1** shows definitions of the datasets, and numbers of mice.

The XGBoost regression model was tested for each of the six available data sets (**Table 1**) using a 10-fold cross-validation approach, with results as shown in **Figure 1**. Each symbol represents an estimate, for each mouse, of the percent increase in lifespan based on the regression equation. The model was run 10 times for each mouse, each time using 90% of the data and omitting 10%. The median estimate for the series of 10 regressions is then plotted in the figure and used to calculate the mean estimated lifespan value (and standard deviation) for the group. Horizontal lines in **Figure 1** indicate these group-specific mean values, and asterisks indicate the outcome of t-tests comparing groups of treated mice to the group of untreated controls (without adjustment for multiple comparisons). **Supplemental Table 1** collects the p-values corresponding to each comparison shown in Figures 1 – 4.

**Figure 1.**
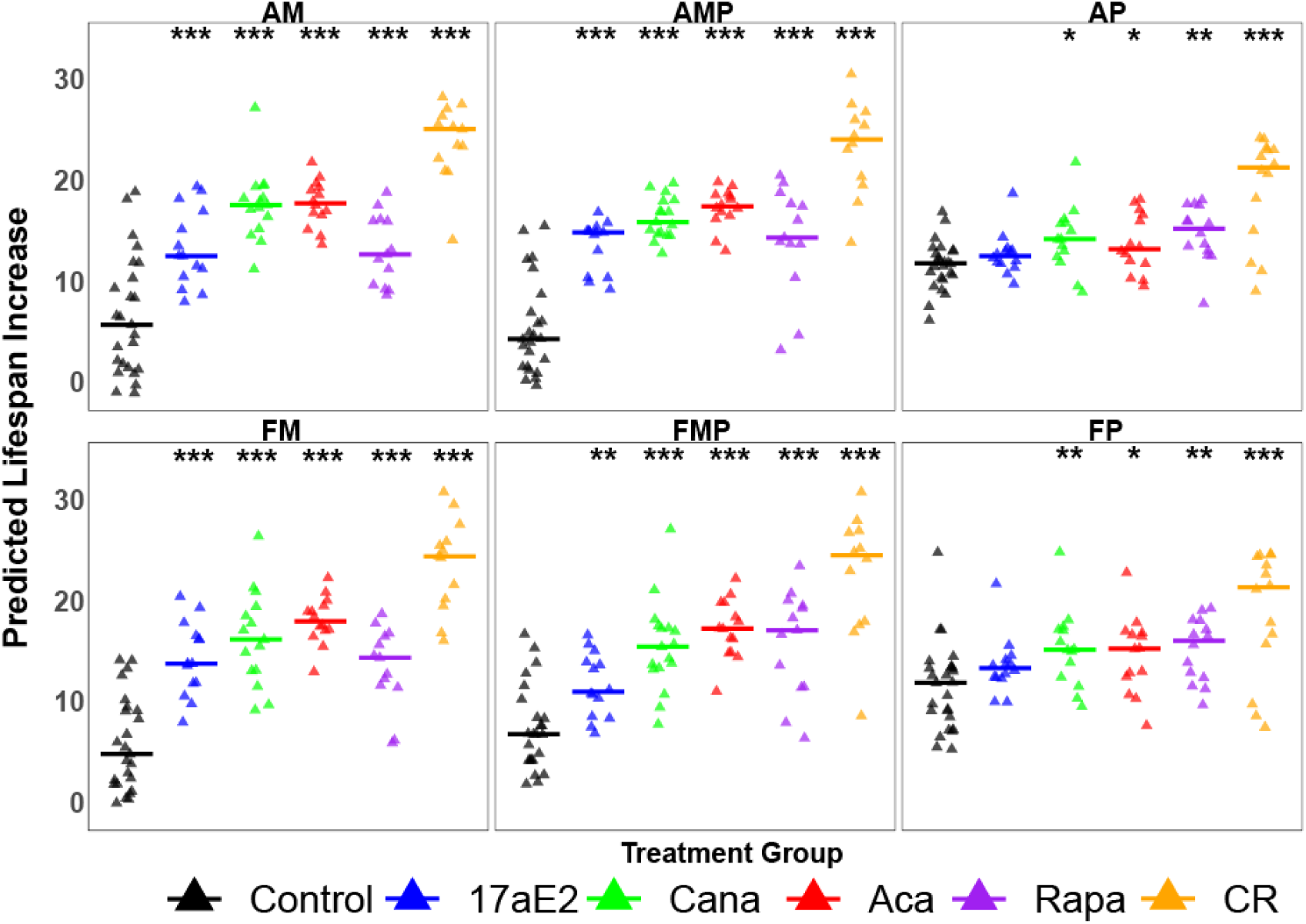
XGBoost regression models including metabolomic features predict percent lifespan increase for male mice for each of five interventions. All data came from analysis of plasma of male mice. Each of the six panels shows results using a different dataset (AM, FM, etc.). Each symbol represents a mouse from the Control group (black symbols) or one of the five indicated intervention groups, evaluated using an XGBoost regression model trained on the indicated data type using 10-fold cross validation (see Methods). The median value for each intervention group is shown as a horizontal line. Asterisks show significance for t-tests contrasting each group of treated mice to control mice, with levels at p = 0.05, 0.01, and 0.001, comparing each group of mice to the control mice.

Each of the metabolomic datasets (AM and FM) suggested that this XGBoost regression approach could distinguish control mice from mice in each of the five treatment groups, using p < 0.05 as criterion. The datasets pooling metabolomic and peptide data (AMP, FMP) were at least equally successful in their ability to discriminate between plasma of control mice and plasma of mice exposed to an anti-aging intervention. The datasets using peptide data only (AP and FP) did less well by this criterion, suggesting higher estimated lifespan increases only for Aca and CR (and for Cana in the FP data).

### Prediction of estimated lifespan increases for novel treatments

To provide a more realistic and pertinent test of the ability of the XGBoost regression method to characterize novel candidate drugs, i.e., interventions that were not used in the training procedure, we used a “novel intervention test” (NIT) procedure, whose results are shown in **Figure 2**. For each mouse in one of the five treatment groups, the model was trained on a dataset consisting of untreated mice and mice in the other four treatment groups but omitting any data from mice in the same treatment group as the tested mouse. This simulates an experimental situation in which the actual lifespan effect of a novel drug is not known and therefore cannot be used in the training set. The estimated lifespan increase of the control mice in Figure 2 was calculated by 20 iterations of the 10-fold cross-validation method used in Figure 1.

**Figure 2.**
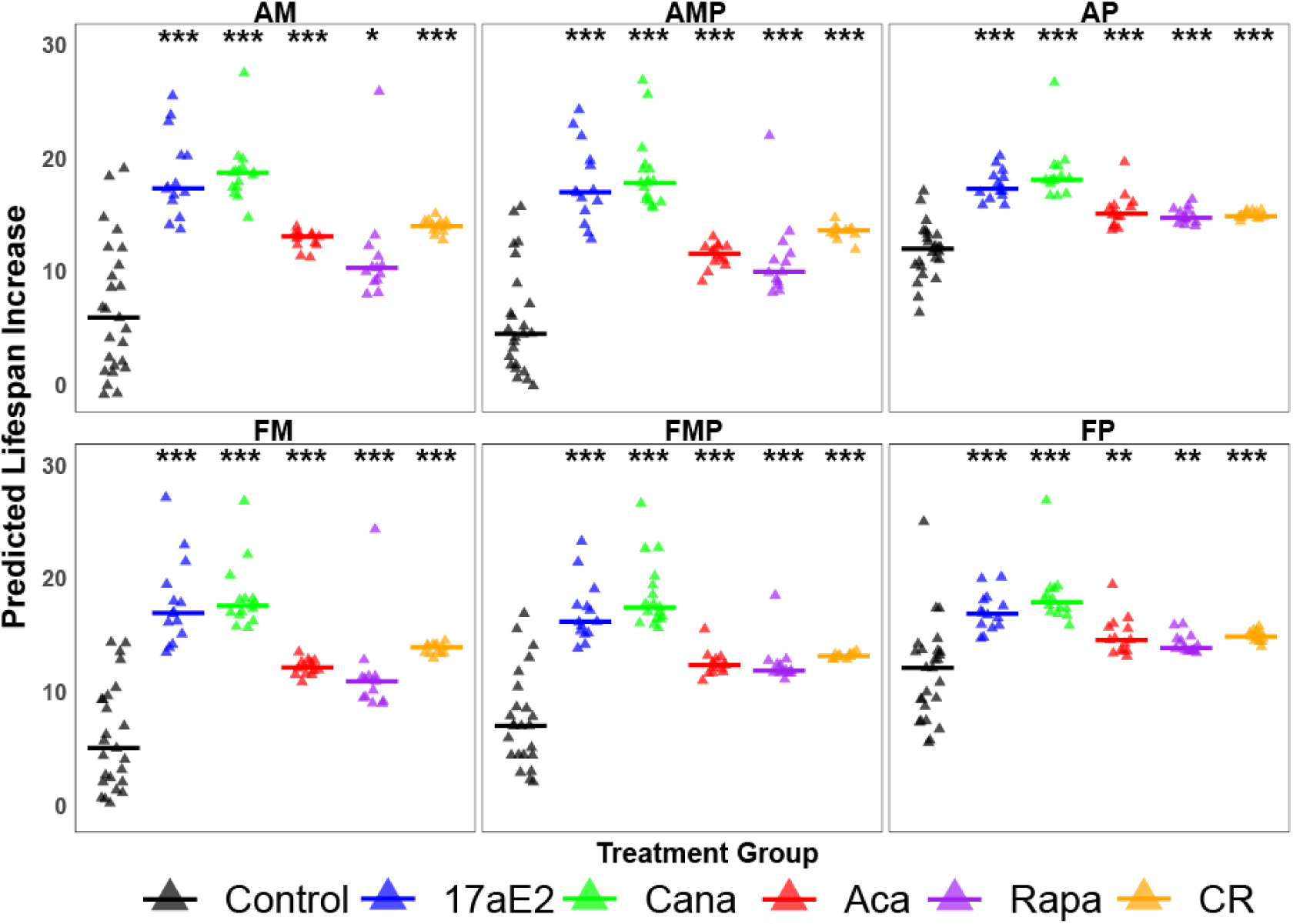
Lifespan estimation for five kinds of slow-aging male mice using XGBoost regression models trained on subsets of four interventions. Each symbol represents one male mouse of the indicated group. For each of the five intervention groups, an XGBoost model trained on controls and the other four treatment groups was used to estimate percent lifespan increase for each mouse in the omitted group (the “novel intervention test.”) For example, symbols for the 17aE2 group reflect a model trained using controls, Cana, CR, Aca, and Rapa mice. Each model was iterated 20 times for each animal, and the median value recorded and plotted in this figure. Horizontal bars show the median value for the group of plotted symbols. Estimated values for control mice were calculated using the same 10-fold cross validation method, trained on the entire set of mice (controls and treated), as used for Figure 1, with 20 iterations for each control mouse. Asterisks show significance for t-tests contrasting each group of treated mice to control mice, with levels at p = 0.05, 0.01, and 0.001. The specific p-values for each group, and other summary information, can be seen in Supplemental Table 1.

The analysis showed that the XGBoost regression model, applied to the NIT design, was able to correctly indicate an elevated estimate of median lifespan increase for each of the five treatments, even when the model was trained on the other four treatments. It is noteworthy that similar performance was achieved even on the datasets (AP and FP) that did not utilize any of the metabolomic data, but relied on peptide data alone.

### Comparison to other ML methods

The results of the XGBoost regression algorithm were compared, via the NIT, to four other ML calculations: support vector regression, K-nearest neighbor regression, HistGradientBoost regression, and random forest regression, using male mice. In each case we applied the ML method to one of the six datasets. **Supplemental Table 2** shows the estimated lifespan increase for each combination of data and method, for each of the five intervention groups, and includes the p-values for a comparison of the treated mice to mice in the untreated control group. Only XGBoost regression was able to distinguish each treated group from controls, for each of the six datasets, with a nominal p < 0.05. The KNN regression method, for example, did not predict significant lifespan increase in 8 of the 30 combinations of data and intervention, and incorrectly estimated a negative effect of Aca for two of the datasets. This comparison is the basis for our use of XGBoost regression for the remainder of the work shown in this paper.

### Lifespan prediction for female mice using XGBoost regression

Development of ML estimation algorithms for male mice was facilitated by the availability of five different interventions, all of which increased male lifespan by at least 12%. We applied the same approach using models trained on data from female mice and found that these were not able to predict percent lifespan increases for either males or females. We suspect that the models failed because only two interventions, Rapa and CR, led to substantial lifespan increase in female mice, although testing this idea will require discovery of additional drugs that extend lifespan in females. To see if a model trained on male data would provide useful insights into metabolomic and/or peptide data obtained from female mice, we calculated the predicted percent lifespan increase for female mice in each of the five treatment classes, using the NIT approach with models trained on male data only. The results are shown in **Figure 3** and showed that models trained using metabolomic data from males (AM, FM, AMP, FMP) predicted lifespan increases when applied to plasma from females treated with 17aE2, Cana, or Aca, or exposed to the CR diet. In contrast, females treated with Rapa would not have been correctly identified as long-lived using a model trained on males treated with the other four interventions, and control males. Models trained using peptide data only (AP or FP) implied lifespan extension for CR females, consistent with the ability of CR diets to extend lifespan in UM-HET3 females. Male-trained models did not predict lifespan extension in mice treated with Aca, Cana, or 17aE2, consistent with the absence of lifespan extension in female mice exposed to Cana or 17aE2, and with the small (though significant) lifespan extension in Aca-treated female mice. Peptide-trained models did not predict a lifespan effect for Rapa-treated females, although Rapa does indeed extend female lifespan. Thus, models developed using data from plasma of five varieties of slow-aging males show mixed success in predicting lifespan benefits based on plasma data from female mice.

**Figure 3.**
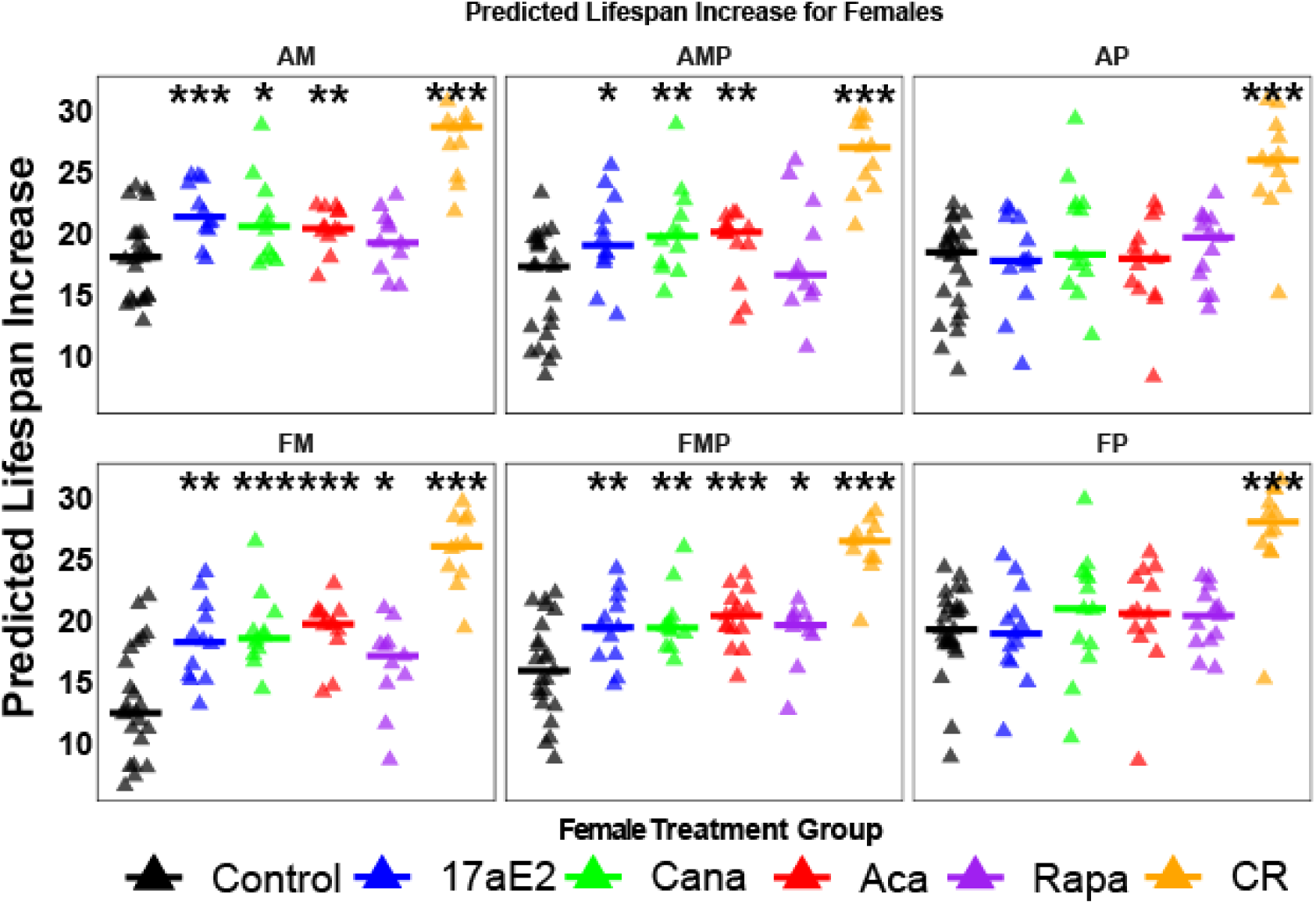
Lifespan estimations for female treated with anti-aging drugs from a male-only trained model. Each symbol represents one female mouse of the indicated group. Each prediction was made once for each mouse. Horizontal bars show the median value for the group of plotted symbols. Asterisks show significance for t-tests contrasting each group of treated mice to control mice, with levels at p = 0.05, 0.01, and 0.001. 17aE2 and Cana are not known to increase lifespan in females. Acarbose, Rapa, and CR are known to increase lifespan in females.

The male-trained models also predicted variable degrees of lifespan extension in untreated female control mice, ranging from 14% - 18%. Female mice of the UM-HET3 stock do typically show lifespans slightly longer than those of male control mice, but the difference is approximately 4% averaged over 13 consecutive annual cohorts.

### Comparison to plasma samples from long-lived mutant mice on two other genetic backgrounds

We also evaluated whether models trained on data from drug- or diet-treated long-lived UM-HET3 mice would reveal similarities to plasma samples from mice carrying mutations that lead to lifespan extension, such as Snell Dwarf (SD) and growth-hormone receptor knockout (GHKRO) mice. The results of these calculations are shown in **Figure 4**. Each of the six datasets (metabolites, peptides, or their combination) predicted higher lifespan for SD mice than for their littermate controls. Similarly, five of the six datasets (FM is the exception) predicted significantly higher lifespan in male GHRKO mice than in their own littermate controls. These results suggest that there is some degree of overlap, in both metabolomic and proteomic plasma changes, between long-lived young adult mutant mice and young adult UM-HET3 mice exposed to anti-aging diets or drugs. It also implies that the usefulness of the XGBoost regression model, in the context of the NIT, is not limited to the UM-HET3 stock alone.

**Figure 4.**
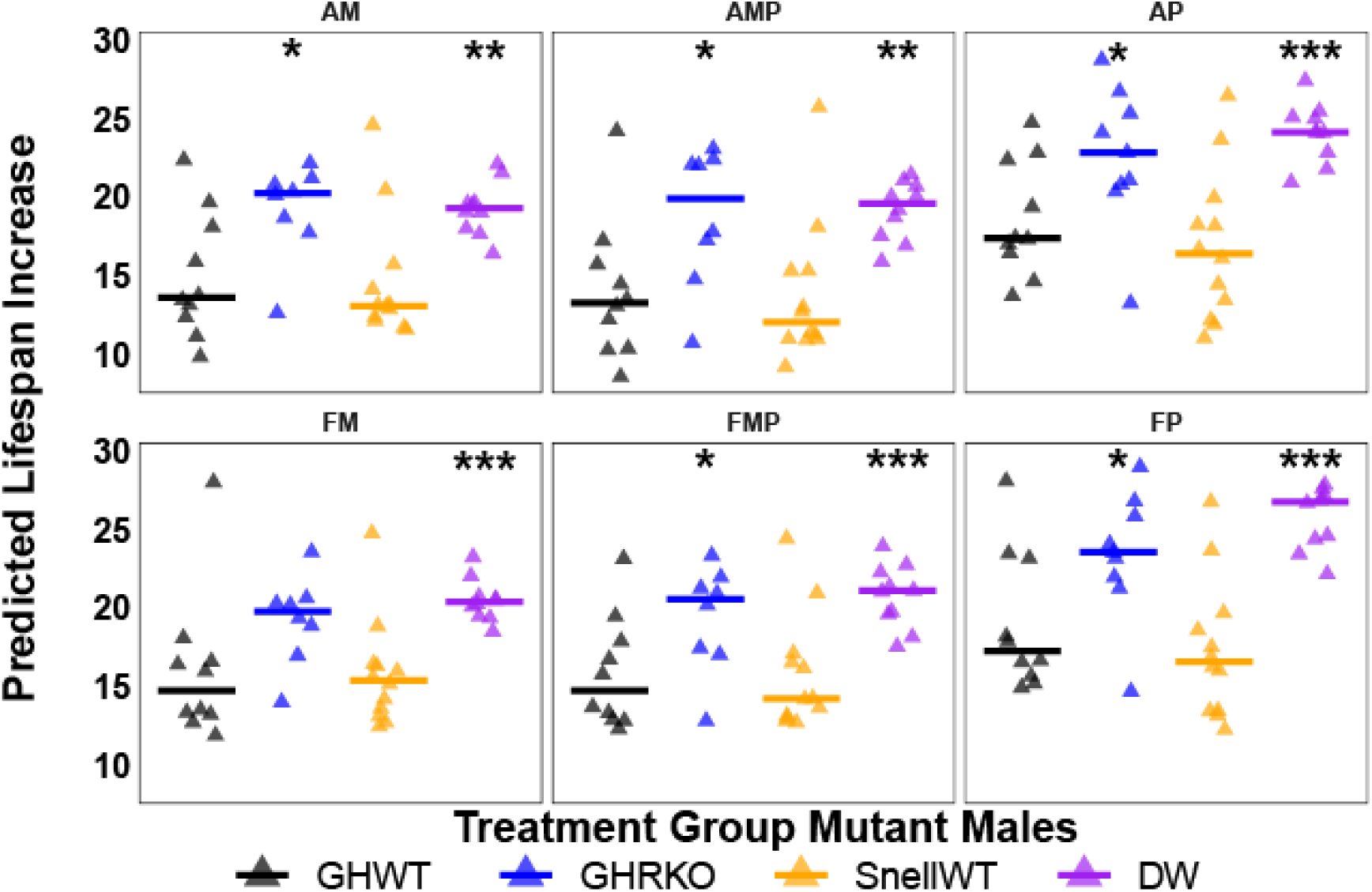
Lifespan estimations for long lived male mutant mice. Each symbol represents one mutant mouse of the indicated group. Each prediction was made once for each mouse. Horizontal bars show the median value for the group of plotted symbols. Asterisks show significance for t-tests contrasting each group of treated mice to control mice, with levels at p = 0.05, 0.01, and 0.001.

### Influential features discriminating control from slow-aging UM-HET3 males

To gain insight into the metabolic features that have the most influence on predicted lifespan change in our UM-HET3 male datasets, we calculated the SHAP value for each metabolite. **Table 3** lists the 20 highest-ranked metabolites based on this criterion, and **Supplemental Table 3** lists the 100 highest-ranked metabolites in our FM-based models.

**Table 3:** Metabolite name and SHAP value rank.

| Rank | Feature Name | Rank | Feature Name |
| --- | --- | --- | --- |
| **1** | N-Methylisoleucine | **11** | N-Acetyltyrosine |
| **2\*** | TG 57:9\|TG 17:1_18:2_22:6 | **12\*** | TG 64:16\|TG 20:4_22:6_22:6 |
| **3\*** | TG 48:3\|TG 14:0_16:1_18:2 | **13\*** | TG O-53:9\|TG O-19:5_17:2_17:2 |
| **4\*** | TG 56:9\|TG 16:1_18:2_22:6 | **14** | N-Acetylaspartylglutamic acid |
| **5** | N-omega-Acetylhistamine | **15** | SM 42:1;2O |
| **6** | Ergothioneine | **16** | Sulfamethoxazole |
| **7** | 1-(Piperidin-4-yl)ethan-1-ol | **17\*** | TG 58:8\|TG 18:1_18:2_22:5 |
| **8** | 3-Hydroxysebacic acid | **18** | N8-Acetylspermidine |
| **9\*** | TG 62:14\|TG 18:2_22:6_22:6 | **19** | SM 36:1;2O\|SM 18:1;2O/18:0 |
| **10\*** | TG 50:3\|TG 16:0_16:1_18:2 | **20** | 1-Hexadecanoyl-2-octadecadienoyl-sn-glycero-3-phosphocholine |

Figure 5 shows the effect of each of the five interventions on each of these 20 features, keyed to the feature number shown in Table 3. Some features (examples: 3, 10) are diminished by all five interventions, with respect to controls. Some features (examples: 9, 12, 18) are increased by all five interventions. There are, however, many features where the direction of the effect is not uniform across all interventions (examples: 2, 4, 5, 8); these features would have been deemed unhelpful using methods that considered only patterns of metabolite change that were uniform in direction across all interventions.

**Figure 5.**
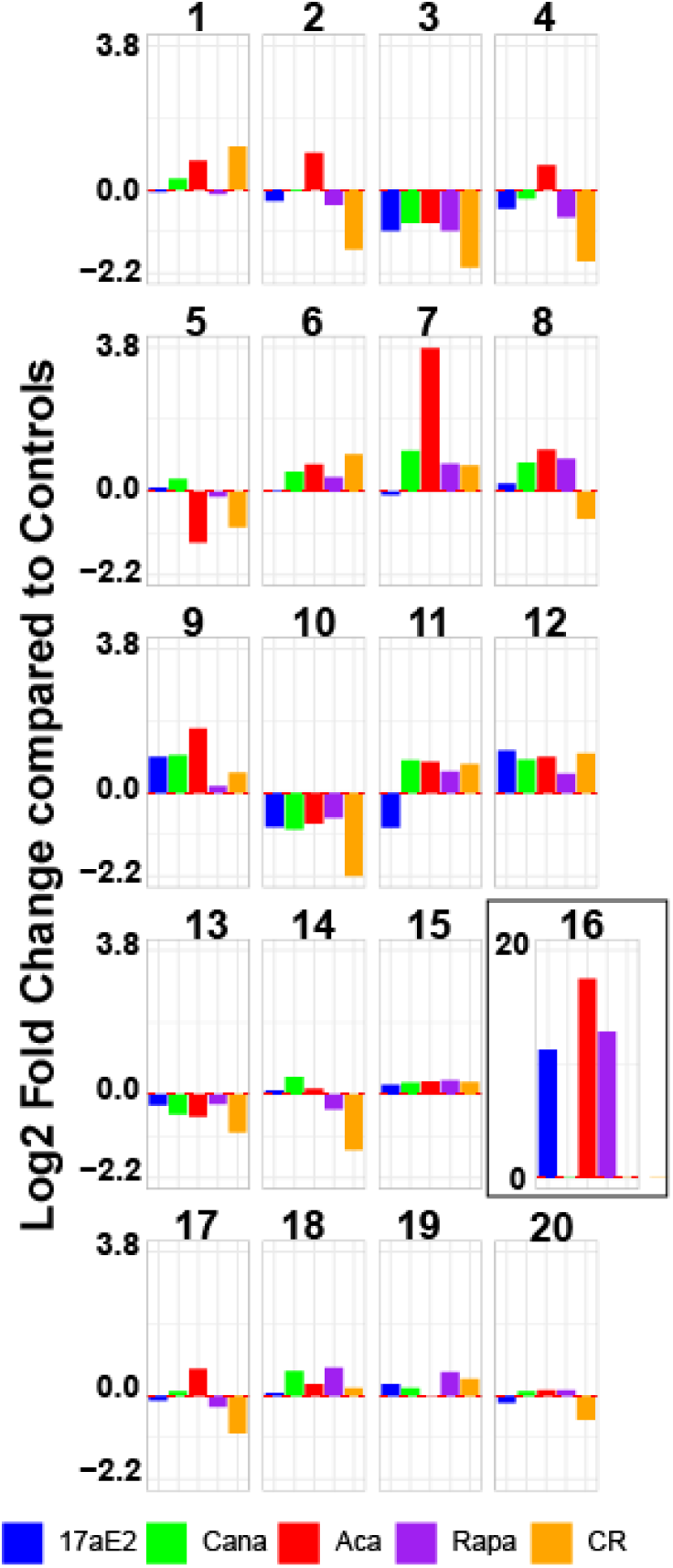
Log2 fold changes of the 20 most informative metabolic features in male mice treated with anti-aging drugs or exposed to the CR diet. Each bar corresponds to the average log2 fold change difference between the treated mice and untreated mice (treated/untreated). The numbers above correspond to the key shown in Table 3. Note altered Y-axis scale for Compound 16; this metabolite was undetectably low in plasma samples from all control mice, and the missing values were therefore replaced by the lowest level measured in treated mice.

Figure 6 displays the “SHAP plots” for each of these 20 metabolic features, indexed to the features shown in Table 3. Each symbol represents one mouse. Red symbols indicate mice in which high values of the metabolite were influential in the contribution of that metabolite to the overall estimated lifespan outcome. Blue symbols show mice in which low values were influential, and purple values show all other mice. Symbols plotted towards the right (values above zero on the X-axis) are mice where the metabolite value implied a higher estimated lifespan, and symbols plotted below X = 0 contributed to lower estimated lifespan.

**Figure 6.**
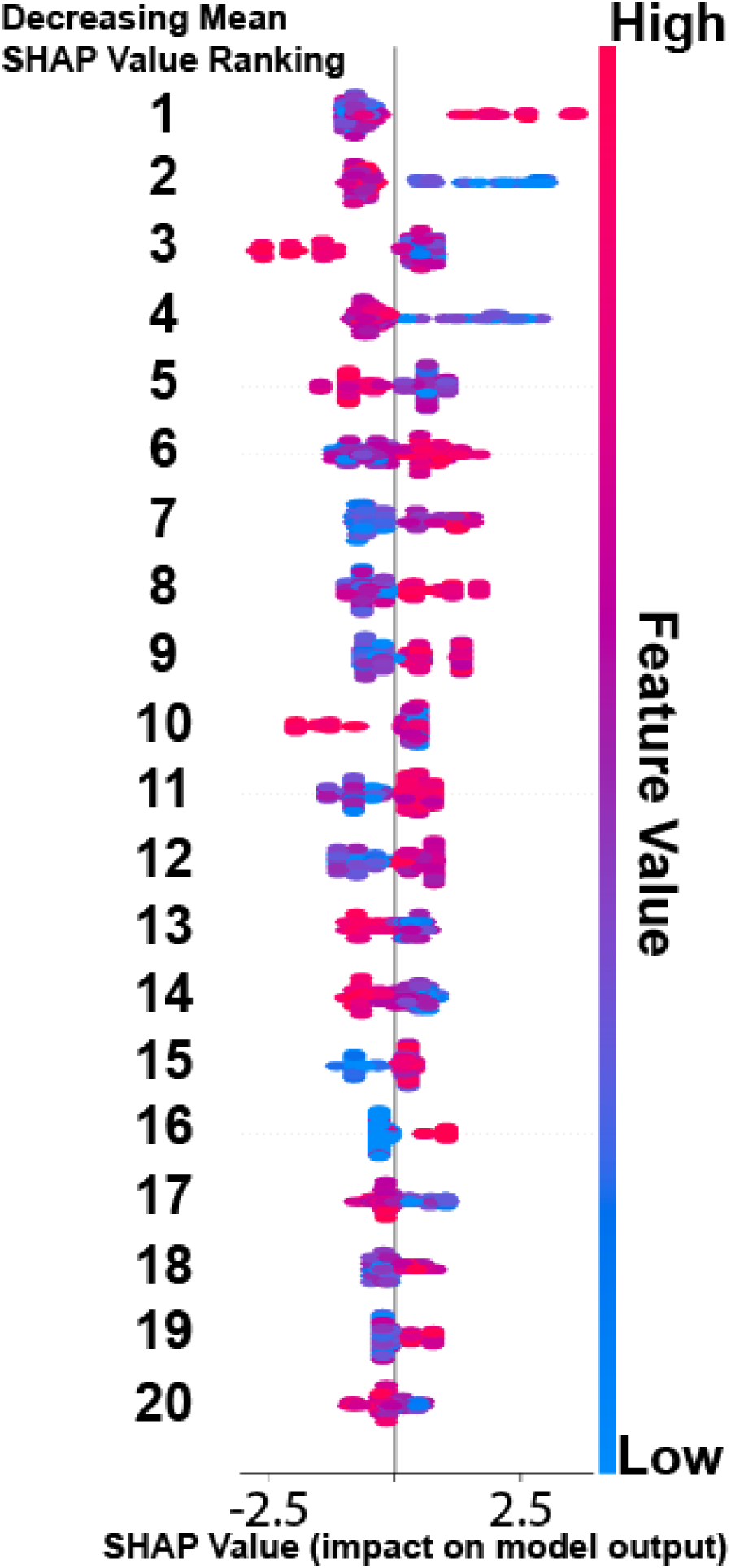
SHAP values for the top 20 predictive features of the model. The color of the SHAP plot corresponds with the magnitude of the feature. Features (1–20) are keyed to the identities listed in Table 3 and shown in Figure 5. Symbols plotted to the right of the vertical line at X = 0 indicate that the value of the feature suggested higher predicted lifespan increase, in an amount proportional to the position of the symbol on the X-axis (SHAP value). Red symbols indicate higher abundance levels, and blue symbols indicate lower abundance.

Of the 20 highest-ranked metabolic features shown in Table 3, 8 are triacylglycerides (TG), suggesting that the anti-aging interventions may often lead to alterations of plasma lipid components. To seek patterns within the TG class, we plotted fatty acid (FA) chain length against double bond number for the 60 FA chains within the 20 highest ranked lipids in the FM data set, using the SHAP score for ranking. Figure 7 shows a clear pattern: with few exceptions, lipids whose FA constituents had longer chain length tended to increase in slow-aging mice, while lipids whose FA chain lengths were shorter (18 carbons or less) tended to decrease in the slow-aging mice.

**Figure 7.**
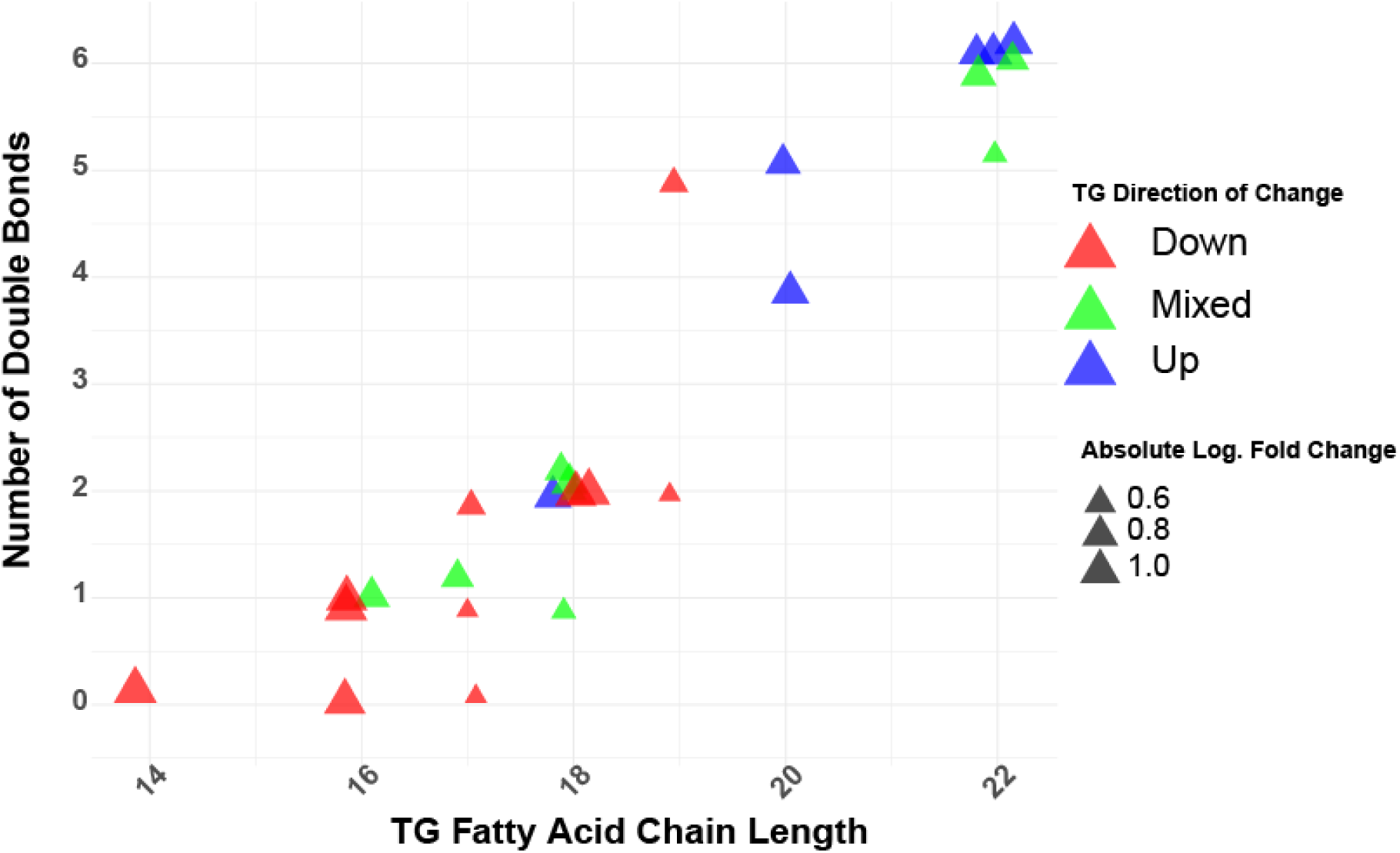
The fatty acid composition of the top 20 most important triacylglycerides in the prediction model. The colors correspond to a change between all treated groups and the untreated control mice. If all the treated groups had lower relative values compared to control mice, then it was colored red; if all the treated groups had higher relative values compared to control mice, then the symbol was colored blue. Green symbols indicate fatty acids which showed a mixture of both increased and decreased levels relative to control mice. The larger the symbol, the larger the magnitude of change in the treated mice compared to the control mice.

## Discussion

In this study, we developed and evaluated a machine learning model using Extreme Gradient Boosting (XGBoost) regression to predict the percentage increase in lifespan of male UM-HET3 mice treated with various five different anti-aging interventions. Using 10-fold cross-validation, in which models are evaluated by successive reservation of 10% of the samples, we found that models that included metabolomic data (AM, FM, AMP, FMP) could successfully differentiate between control mice and mice exposed to any of five interventions that extend lifespan of males. Tests on datasets containing only peptide data were slightly less successful; in particular, they did not predict higher lifespan in mice treated with 17aE2.

The “novel intervention test” (NIT) provides a better simulation of a situation in which a candidate drug, not previously tested for lifespan effect, would be evaluated using XGBoost regression trained on data from validated anti-aging drugs and sex-matched controls. In our setting, the model is trained on four of the validated interventions, and estimated percent lifespan increases are calculated for each mouse in the group excluded from the training set. Lifespan estimates are then compared to those of untreated control mice, estimated by the 10-fold cross-validation calculation. As shown in Figure 2, each of the five varieties of treated male mice produced an estimated mean level of lifespan increase that is significantly higher than that of control animals, for datasets comprising metabolites, peptides, or a combination of both kinds of data.

It is noteworthy that all the mice tested in this study were 12 months of age when plasma was taken for testing. This design feature helps minimize effects of aging and age-related diseases on plasma constituents. It also more closely mimics a design in which new candidate drugs could be evaluated by administration to young adults for a comparatively short interval, in the current case 8 months, at less expense and in shorter time than needed for a complete lifespan study. One goal for future work is to determine if this approach can help to prioritize candidate drugs to select a subset that deserves full-scale lifespan testing.

We were unable to develop successful XGBoost models using data from female mice only, and we suspect that this is because only two of the interventions, CR and Rapa, produce lifespan benefits that exceed 7%. We found, however, that the metabolite-trained models developed using FM or FMP data from male mice would have implied extended lifespan in females exposed to any of the five anti-aging interventions (see Figure 3). This implies that some of the metabolic changes induced by anti-aging interventions in male mice are also produced in female mice, even for drugs (17aE2, Cana) that do not produce significant increases in female lifespan at the doses use. It is possible that these agents might produce a combination of beneficial effects in female mice that are obscured by harmful side-effects that limit lifespan (26). Further exploration of sex effects in this system may have to await discovery of other interventions that extend lifespan of female mice.

To see if the metabolic features induced by drugs and diet were also characteristic of mice whose lifespan had been lengthened by a genetic mutation, we looked at plasma from 12 month old males of two such stocks: Snell dwarf, which are deficient in growth hormone, thyroid stimulating hormone, and prolactin, and GHRKO mice, whose extended longevity is thought to be related to deficient responses to GH alone, without hypothyroidism or low prolactin (14, 27–29). We found that all six models (trained on metabolites, peptides, or both) predicted elevated lifespans for Snell Dwarf mice, and that five of the datasets made similar correct predictions for lifespan increase in GHRKO mice. The result implies some similarities in metabolic and proteomic effects shared by both mutant and drug/diet-treated mice, and that the success of the XGBoost/NIT procedure is not limited to the UM-HET3 genetic stock. Tests of additional long-lived mutants, such as PAPPA-KO and PTEN-transgenic overexpressors, will be needed to explore this issue further.

Of the 20 metabolites that had the greatest influence on the estimation of lifespan predictions (FM dataset), 8/20 (40%) were triacylglycerides (TG), although TG made up only 16% of the metabolites in this dataset (p = 0.003). Among the 20 highest-ranked TG (see **Supplemental Table 2**), FA with 20 – 22 carbon chains were most often at higher levels in the long-lived mice, while FA with 14 – 18 carbon chains were typically down-regulated. Interestingly, studies in model organisms have demonstrated that manipulation of lipid metabolism can influence lifespan. For example, overexpression of fatty acid desaturase genes, leading to increased production of unsaturated fatty acids like DHA, has been associated with extended lifespan in nematodes (30).

To facilitate the interpretation of the many metabolites in the FM dataset, we developed a freely accessible web tool that displays the relative concentrations of all detected metabolites in both male and female mice from this study, as well as in 6-month-old “Young” UM-HET3 mice. This user-friendly visualization tool enables both researchers and non-experts to explore whether, and how, specific metabolites of interest differ in control versus long-lived mice. Statistical significance (p < 0.05), determined by Welch’s two-tailed t-test, is indicated by a red asterisk, highlighting differences relative to same-sex controls. The details on where to access the tool can be found in the methods section.

Previous work from this lab (31) has documented shared features seen in each of 10 different varieties of slow aging mice, including the 7 kinds of mice used in the present paper, and argued that these can serve as aging rate indicators (ARI). Conceptually, ARIs are measures of the instantaneous rate of aging, i.e., the rate at which aging is occurring at the time of sample collection. Unlike biomarkers, ARIs do not require assessment at old age, or at two consecutive ages, since they do not depend on age (or “biological age”), but instead report the rate of age-related change. ARIs shown to date include changes in fat, macrophages, muscle, liver, and brain, as well as two (GPLD1, irisin) proteins present in mouse plasma. We speculate that some of the features shown in Figures 5 and 6 may prove useful as plasma-based ARIs in future studies and could be put to use in screening of candidate anti-aging drugs in mice, dogs, and/or humans.

The present work has many limitations, which can be addressed by future studies. For one thing, we did not have access to plasma from 12-month-old mice treated with drugs known not to increase UM-HET3 lifespan, and will need to acquire new samples of this kind to see if the NIT can successfully discriminate drugs known to extend mouse lifespan from those known not to do so. Next, it will be helpful to test the robustness of this approach to data on slow-aging mice generated by other laboratories, in other background stocks, and using other metabolomic pipelines. Application of the approach to metabolic data obtained from dogs of breeds or body weights that differ in expected lifespan will also be of interest. It will be useful to learn if the XGBoost/NIT approach has similar success when applied to internal tissues (liver, brain, kidney, etc.) of slow-aging mice, and, if so, to see to what extent the most influential features in the plasma dataset might overlap with similar lists prepared from various internal tissues. It is also possible that some of these metabolic and proteomic features, in combination, could provide useful insights into changes induced by short-term administration of potential anti-aging drugs in human volunteers.

In summary, our XGBoost regression models suggest that there may be shared metabolic changes produced, at least in male mice, by CR diets and each of four anti-aging drugs with varying biochemical targets and mechanisms. These changes can be seen in 12-month-old (young adult) mice after 8 months of treatment. Some of the changes are also seen in female mice exposed to the same set of treatments as young adults. The alterations can be detected in plasma, which may provide a useful bridge to studies of humans, in which plasma samples are more readily obtainable than samples of internal tissues. Some of the metabolites that have particularly strong influence on the calculated longevity estimates may be useful as aging rate indicators and help to prioritize new candidate drugs for more intensive evaluation. The list of metabolites with high SHAP scores in our study also implicate specific metabolic and physiological pathways that might help control aging rate and might usefully be evaluated as targets for discovery of new anti-aging drugs or nutritional interventions.

## Acknowledgements

We are grateful to Lori Roberts, Jacob Sheets, Micah Bush, Lindsey Burger, and Robert Dilg for expert assistance in mouse husbandry and specimen preparation. This work was supported by NIH grants AG023122, AG024824, T32-AG000114 and R35 GM137795.

## Acknowledgements

We thank Dr. Robert Moritz for providing the plasma peptide data used in this paper.

## Supplemental Materials

**Supplemental Table 1:** compilation of statistical results for contrasts shown in Figures 1 – 4.

| Figure # | Dataset | Sex_Treatment | Pval |
| --- | --- | --- | --- |
| 1 | AM | M_17aE2 | 8.1E-07 |
| 1 | AM | M_Aca | 2.5E-12 |
| 1 | AM | M_CR | 4.0E-12 |
| 1 | AM | M_Cana | 1.6E-08 |
| 1 | AM | M_Rapa | 6.8E-05 |
| 1 | AMP | M_17aE2 | 9.1E-08 |
| 1 | AMP | M_Aca | 4.3E-12 |
| 1 | AMP | M_CR | 1.2E-11 |
| 1 | AMP | M_Cana | 2.0E-10 |
| 1 | AMP | M_Rapa | 2.1E-06 |
| 1 | AP | M_17aE2 | 1.9E-01 |
| 1 | AP | M_Aca | 3.9E-02 |
| 1 | AP | M_CR | 4.1E-05 |
| 1 | AP | M_Cana | 1.6E-03 |
| 1 | AP | M_Rapa | 9.5E-04 |
| 1 | FM | M_17aE2 | 2.7E-06 |
| 1 | FM | M_Aca | 3.0E-13 |
| 1 | FM | M_CR | 1.2E-11 |
| 1 | FM | M_Cana | 5.5E-08 |
| 1 | FM | M_Rapa | 6.2E-07 |
| 1 | FMP | M_17aE2 | 3.7E-06 |
| 1 | FMP | M_Aca | 2.1E-13 |
| 1 | FMP | M_CR | 1.0E-08 |
| 1 | FMP | M_Cana | 1.4E-11 |
| 1 | FMP | M_Rapa | 1.6E-07 |
| 1 | FP | M_17aE2 | 8.6E-02 |
| 1 | FP | M_Aca | 3.9E-03 |
| 1 | FP | M_CR | 9.1E-04 |
| 1 | FP | M_Cana | 2.1E-03 |
| 1 | FP | M_Rapa | 2.8E-04 |
| 2 | AM | M_17aE2 | 4.3E-09 |
| 2 | AM | M_Aca | 2.6E-05 |
| 2 | AM | M_CR | 2.0E-06 |
| 2 | AM | M_Cana | 1.6E-10 |
| 2 | AM | M_Rapa | 1.1E-02 |
| 2 | AMP | M_17aE2 | 2.1E-10 |
| 2 | AMP | M_Aca | 4.5E-06 |
| 2 | AMP | M_CR | 2.9E-08 |
| 2 | AMP | M_Cana | 7.3E-13 |
| 2 | AMP | M_Rapa | 4.4E-04 |
| 2 | AP | M_17aE2 | 3.7E-12 |
| 2 | AP | M_Aca | 1.6E-06 |
| 2 | AP | M_CR | 3.4E-07 |
| 2 | AP | M_Cana | 5.9E-09 |
| 2 | AP | M_Rapa | 5.9E-07 |
| 2 | FM | M_17aE2 | 9.9E-10 |
| 2 | FM | M_Aca | 6.8E-07 |
| 2 | FM | M_CR | 1.2E-08 |
| 2 | FM | M_Cana | 1.2E-12 |
| 2 | FM | M_Rapa | 7.8E-04 |
| 2 | FMP | M_17aE2 | 5.2E-10 |
| 2 | FMP | M_Aca | 8.3E-06 |
| 2 | FMP | M_CR | 1.1E-06 |
| 2 | FMP | M_Cana | 5.3E-12 |
| 2 | FMP | M_Rapa | 2.2E-05 |
| 2 | FP | M_17aE2 | 2.4E-06 |
| 2 | FP | M_Aca | 1.7E-03 |
| 2 | FP | M_CR | 8.3E-04 |
| 2 | FP | M_Cana | 3.6E-07 |
| 2 | FP | M_Rapa | 6.7E-03 |
| 3 | AM | F_17aE2 | 6.2E-04 |
| 3 | AM | F_Aca | 5.5E-03 |
| 3 | AM | F_CR | 1.0E-08 |
| 3 | AM | F_Cana | 2.4E-02 |
| 3 | AM | F_Rapa | 1.8E-01 |
| 3 | AMP | F_17aE2 | 1.3E-02 |
| 3 | AMP | F_Aca | 6.1E-03 |
| 3 | AMP | F_CR | 6.9E-09 |
| 3 | AMP | F_Cana | 4.5E-03 |
| 3 | AMP | F_Rapa | 2.1E-01 |
| 3 | AP | F_17aE2 | 7.0E-01 |
| 3 | AP | F_Aca | 8.5E-01 |
| 3 | AP | F_CR | 1.2E-05 |
| 3 | AP | F_Cana | 1.2E-01 |
| 3 | AP | F_Rapa | 1.8E-01 |
| 3 | FM | F_17aE2 | 1.2E-03 |
| 3 | FM | F_Aca | 1.1E-05 |
| 3 | FM | F_CR | 2.7E-10 |
| 3 | FM | F_Cana | 1.2E-04 |
| 3 | FM | F_Rapa | 5.2E-02 |
| 3 | FMP | F_17aE2 | 1.0E-02 |
| 3 | FMP | F_Aca | 2.7E-04 |
| 3 | FMP | F_CR | 6.3E-10 |
| 3 | FMP | F_Cana | 1.7E-03 |
| 3 | FMP | F_Rapa | 1.5E-02 |
| 3 | FP | F_17aE2 | 8.8E-01 |
| 3 | FP | F_Aca | 3.8E-01 |
| 3 | FP | F_CR | 2.1E-05 |
| 3 | FP | F_Cana | 3.6E-01 |
| 3 | FP | F_Rapa | 3.2E-01 |
| 4 | AM | M_DW | 2.1E-03 |
| 4 | AM | M_GHRKO | 2.0E-02 |
| 4 | AMP | M_DW | 1.8E-03 |
| 4 | AMP | M_GHRKO | 3.8E-02 |
| 4 | AP | M_DW | 2.3E-04 |
| 4 | AP | M_GHRKO | 5.2E-02 |
| 4 | FM | M_DW | 5.0E-04 |
| 4 | FM | M_GHRKO | 8.8E-02 |
| 4 | FMP | M_DW | 7.4E-04 |
| 4 | FMP | M_GHRKO | 4.6E-02 |
| 4 | FP | M_DW | 2.0E-05 |
| 4 | FP | M_GHRKO | 3.7E-02 |

**Supplemental Table 2:** comparison of ML regression models. Each pair of columns presents the results of the Novel Intervention Test applied to data from male mice using the ML regression approach listed in the column header. “Lifespan” values are estimated percentage increase in median lifespan averaged over each mouse in the control or tested group. The “P Value” columns give −log10(p) for the t-test comparison of mice in the indicated treatment group to Control mice. Estimates of Control mouse lifespan are based on the 10-fold cross-validation method illustrated in Figure 1. Red highlights indicate p-values that do not reach the p = 0.05 significance criteria, or lifespan increases that are below zero.

| Dataset | Mouse Group | XGBoost |  | HistGradientBoost |  | KNN |  | Random Forest |  | SVR |  |
| --- | --- | --- | --- | --- | --- | --- | --- | --- | --- | --- | --- |
|  |  | Lifespan | P Value | Lifespan | P Value | Lifespan | P Value | Lifespan | P Value | Lifespan | P Value |
| AM | Control | 7 |  | 5 |  | 10 |  | 7 |  | 7 |  |
| AM | 17aE2 | 19 | 8.4 | 19 | 17.9 | 18 | 3.1 | 18 | 6.7 | 18 | 8.8 |
| AM | Aca | 13 | 4.6 | 4 | 0.4 | 8 | 1.3 | 0 | 7.0 | 4 | 1.7 |
| AM | Cana | 19 | 9.8 | 19 | 10.9 | 13 | 1.4 | 20 | 11.9 | 18 | 11.4 |
| AM | CR | 14 | 5.7 | 14 | 11.8 | 15 | 3.6 | 13 | 5.6 | 14 | 7.4 |
| AM | Rapa | 12 | 2.0 | 9 | 2.4 | 13 | 0.8 | 10 | 0.9 | 12 | 1.6 |
| AMP | Control | 6 |  | 5 |  | 10 |  | 8 |  | 7 |  |
| AMP | 17aE2 | 18 | 9.7 | 22 | 15.7 | 17 | 2.9 | 19 | 7.6 | 18 | 7.7 |
| AMP | Aca | 12 | 5.3 | 4 | 1.1 | 9 | 0.4 | 1 | 6.5 | 6 | 1.1 |
| AMP | Cana | 19 | 12.1 | 20 | 13.7 | 14 | 1.9 | 20 | 10.8 | 18 | 10.4 |
| AMP | CR | 14 | 7.5 | 13 | 9.1 | 13 | 2.2 | 12 | 3.4 | 13 | 6.4 |
| AMP | Rapa | 12 | 3.4 | 10 | 2.5 | 12 | 0.9 | 9 | 0.2 | 11 | 1.2 |
| AP | Control | 12 |  | 9 |  | 11 |  | 10 |  | 11 |  |
| AP | 17aE2 | 18 | 11.4 | 17 | 7.8 | 14 | 1.9 | 14 | 3.6 | 18 | 4.3 |
| AP | Aca | 16 | 5.8 | 4 | 4.8 | 3 | 6.1 | 0 | 16.5 | 0 | 10.9 |
| AP | Cana | 19 | 8.2 | 20 | 8.0 | 13 | 0.6 | 16 | 9.5 | 17 | 4.9 |
| AP | CR | 15 | 6.5 | 7 | 0.5 | 7 | 2.7 | 8 | 2.3 | 10 | 0.6 |
| AP | Rapa | 15 | 6.2 | 6 | 2.4 | 12 | 0.9 | 9 | 0.6 | 7 | 2.0 |
| FM | Control | 6 |  | 4 |  | 5 |  | 6 |  | 5 |  |
| FM | 17aE2 | 18 | 9.0 | 21 | 20.7 | 24 | 14.0 | 23 | 19.2 | 17 | 4.6 |
| FM | Aca | 13 | 6.2 | 3 | 1.0 | 0 | 7.0 | 0 | 8.7 | -3 | 5.0 |
| FM | Cana | 19 | 11.9 | 17 | 8.1 | 7 | 0.3 | 17 | 9.7 | 15 | 4.6 |
| FM | CR | 14 | 7.9 | 12 | 14.1 | 12 | 9.3 | 14 | 10.4 | 14 | 3.6 |
| FM | Rapa | 12 | 3.1 | 8 | 1.8 | 12 | 9.2 | 10 | 1.5 | 18 | 6.0 |
| FMP | Control | 8 |  | 6 |  | 5 |  | 8 |  | 6 |  |
| FMP | 17aE2 | 17 | 9.3 | 21 | 16.9 | 23 | 13.7 | 22 | 13.8 | 18 | 6.8 |
| FMP | Aca | 13 | 5.1 | 3 | 2.4 | 0 | 6.6 | 0 | 8.2 | -1 | 5.9 |
| FMP | Cana | 19 | 11.3 | 18 | 7.5 | 6 | 0.2 | 17 | 6.0 | 16 | 6.1 |
| FMP | CR | 14 | 6.0 | 12 | 7.0 | 12 | 8.0 | 13 | 4.2 | 13 | 4.3 |
| FMP | Rapa | 13 | 4.7 | 10 | 2.4 | 12 | 9.3 | 8 | 0.0 | 13 | 3.6 |
| FP | Control | 12 |  | 8 |  | 4 |  | 9 |  | 7 |  |
| FP | 17aE2 | 17 | 5.6 | 18 | 7.8 | 15 | 9.1 | 15 | 5.8 | 17 | 4.0 |
| FP | Aca | 15 | 2.8 | 3 | 4.3 | 0 | 2.9 | 0 | 15.1 | -2 | 5.7 |
| FP | Cana | 19 | 6.4 | 20 | 9.2 | 15 | 5.2 | 16 | 6.9 | 19 | 6.6 |
| FP | CR | 15 | 3.1 | 8 | 0.1 | 6 | 0.4 | 7 | 1.9 | 11 | 3.0 |
| FP | Rapa | 15 | 2.2 | 7 | 0.7 | 12 | 6.4 | 9 | 0.0 | 1 | 6.0 |

**Supplemental Table 3.**
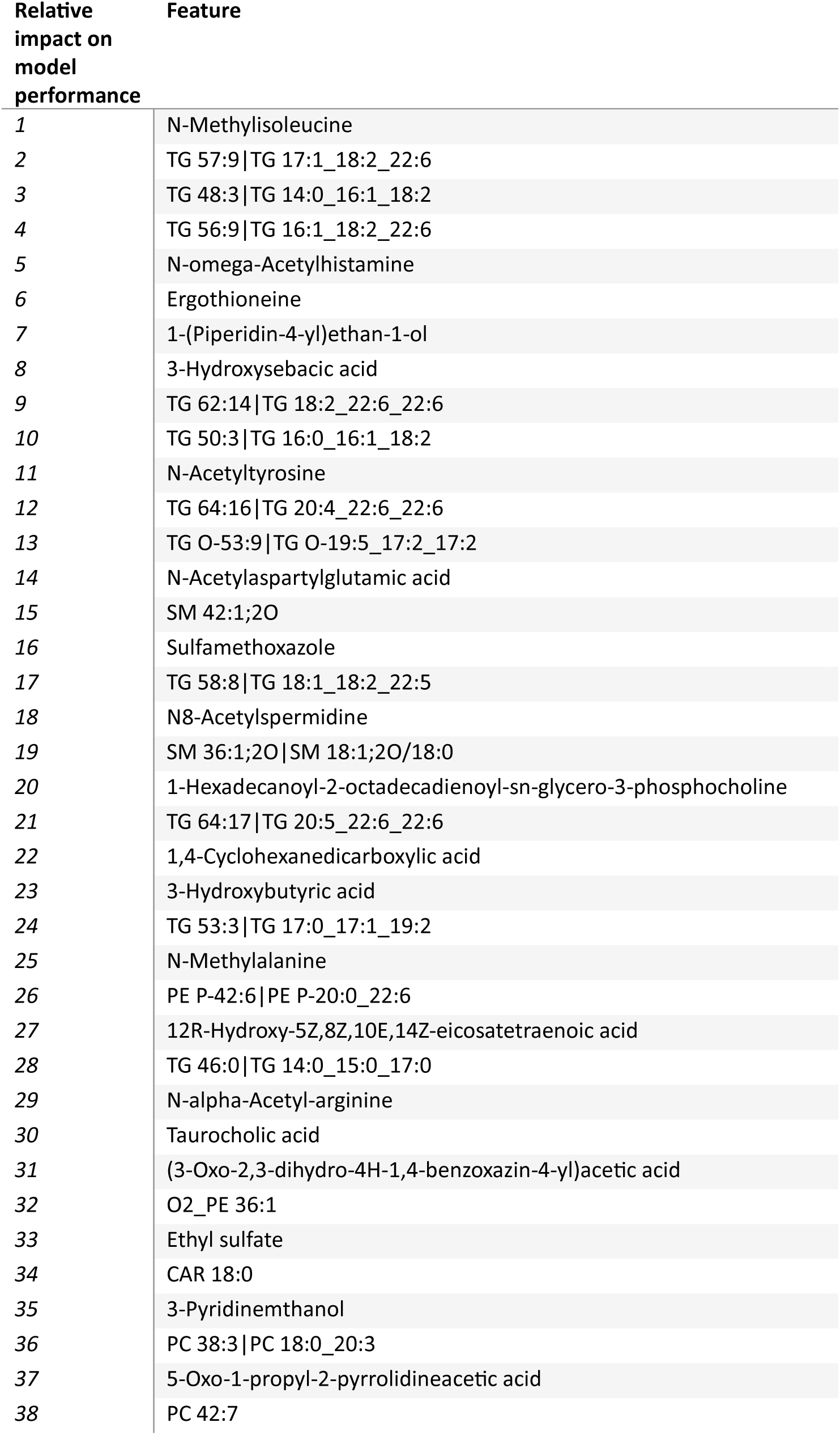
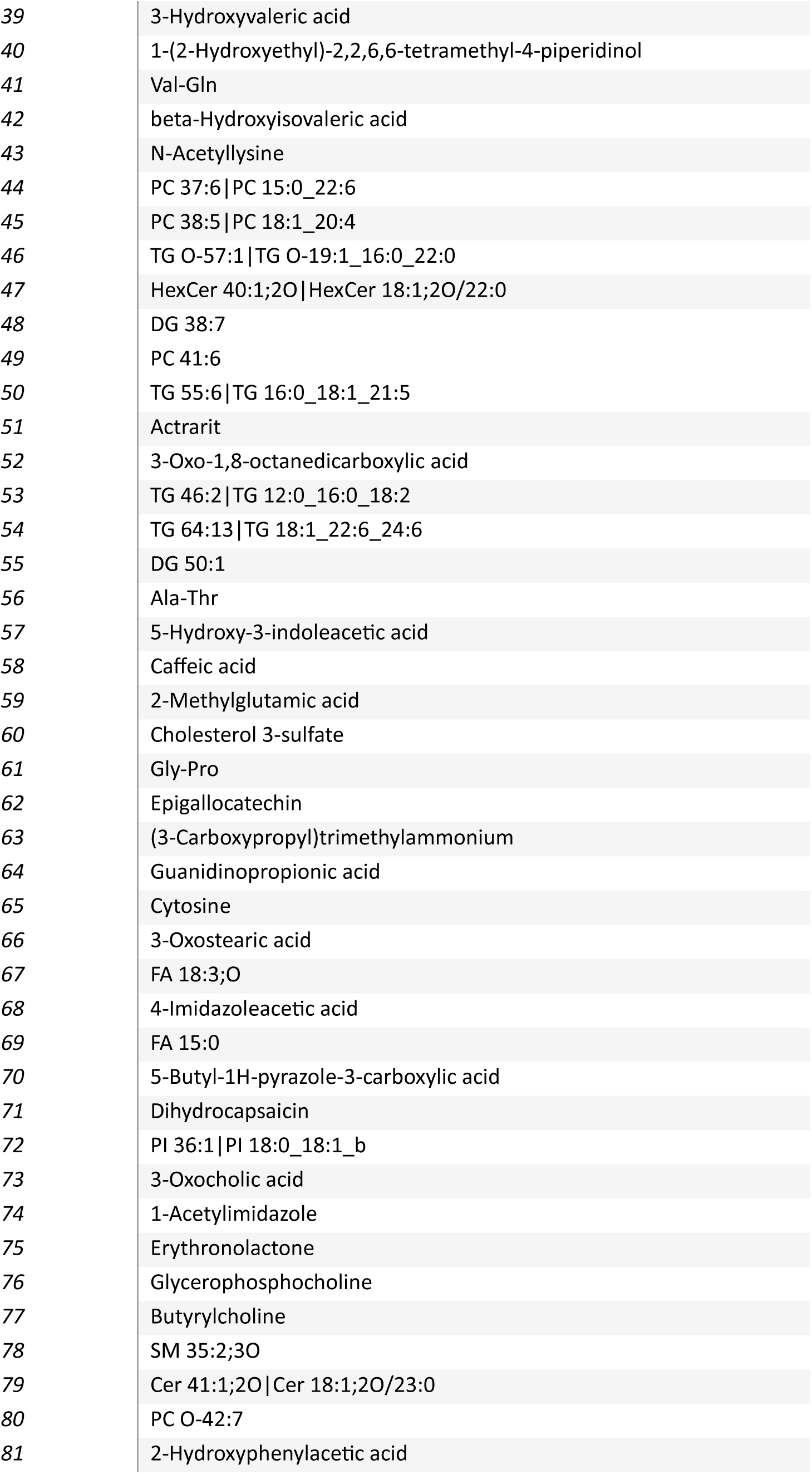
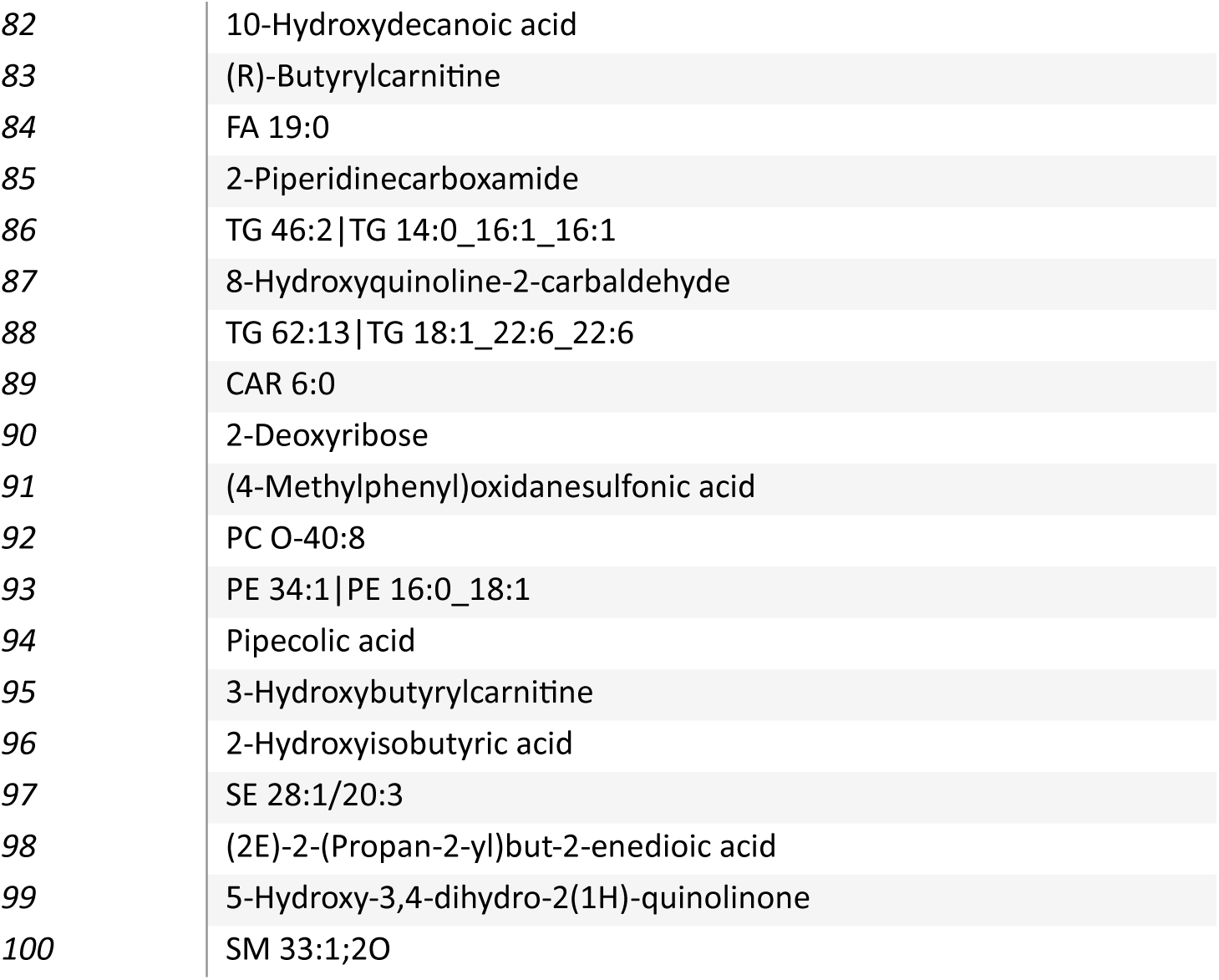
The top 100 ranked SHAP scores after 100 iterations of the XGBoost regression algorithm using the FM dataset.

